# A refined human linear B cell epitope map of Outer surface protein C (OspC) from the Lyme disease spirochete, *Borreliella burgdorferi*

**DOI:** 10.1101/2024.05.29.596441

**Authors:** Grace Freeman-Gallant, Kathleen McCarthy, Jennifer Yates, Karen Kulas, Michael J. Rudolph, David J Vance, Nicholas J Mantis

## Abstract

A detailed understanding of the human antibody response to <u>O</u>uter <u>s</u>urface <u>p</u>rotein C (OspC) of *Borrelellia burgdorferi* has important implications for Lyme disease diagnostics and vaccines. In this report, a total of 13 peptides encompassing eight reported OspC linear B cell epitopes from OspC types A, B and K, including the conserved C-terminus (residues 193-210: peptide C10), were evaluated by multiplex immunoassay (MIA) for IgG reactivity with ∼700 human serum samples confirmed positive in a two-tiered Lyme disease diagnostic assay and ∼160 post-treatment Lyme disease (PTLD) serum samples. The VlsE C6-17 peptide was included as a positive control. Diagnostic serum IgG reacted with 11 of the 13 OspC-derived peptides, significantly more than controls, with the C10 peptide being the most reactive. In the PTLD serum samples, two OspC peptides including C10 were significantly more reactive than controls. Spearman’s rank correlation matrices and hierarchical clustering indicated a strong correlation between C10 and VlsE C6-17 peptide reactivity but little demonstrable association between C10 and the other OspC peptides or recombinant OspC. OspC peptide reactivities (excluding C10) were strongly correlated with each other and were disproportionately influenced by a subset of pan-reactive samples. In the PTLD cohort, C10 clustered with the other OspC-derived peptides and was distinct from OspC and VlsE C6-17. The asynchronous serologic response to OspC, C10, and the OspC-derived peptides reveals the complexity of B cell responses to *B. burgdorferi* and confounds simple interpretation of antibody profiles associated with Lyme disease.

**IMPORTANCE:** Lyme disease is an emerging tick-borne infection caused by the spirochete, *Borreliella burgdorferi*. In humans, antibodies against spirochetal outer surface lipoproteins are proposed to play a role in disease resolution and in protection against reinfection. Some of those same antibodies also serve as diagnostic indicators of an active or history of Lyme disease. In this study, we sought to validate reported antibody binding sites on Outer surface protein C (OspC), a known target of both protective and diagnostic antibodies.

## INTRODUCTION

Lyme (borreliosis) disease is the most common vector-borne infection in the United States, with an estimated 450,000 cases per year (1). The primary etiologic agent of Lyme disease is the spirochetal bacterium, *Borreliella burgdorferi*. In North America, the spirochete is transmitted to humans by black legged ticks, *Ixodes scapularis* and *Ixodes pacificus*, during the course of a blood meal. The spirochete proliferates at the site of the tick bite, typically resulting in an expanding skin lesion commonly referred to as erythema migrans (2-4). In the absence of antibiotic intervention, *B. burgdorferi* can disseminate to peripheral tissues, organs, joints, and the central nervous system, potentially resulting in complications including neuroborreliosis, carditis and/or Lyme arthritis weeks, months or even years later (2, 5). Moreover, a small fraction of Lyme disease patients who receive a full regimen of antibiotics will report persistent health issues (e.g., fatigue, cognitive issues, musculoskeletal pain), a difficult to define syndrome termed post-treatment Lyme disease (PTLD) (6-9). Developing rapid and accurate diagnostic tests capable of detecting and discriminating between LD, PTLD syndrome and *B. burgdorferi* reinfection are much needed (21, 10).

The humoral response to *B. burgdorferi* is robust, resulting in detectable spirochete-specific B cells and serum IgM and IgG days and weeks following an infectious tick bite (11-14). In Lyme disease patients, antibodies are primarily directed against *B. burgdorferi*’s panoply of outer surface lipoproteins (15). Among these, Outer surface protein C (OspC; also known as BB_B19, P23 and P25) stands out. OspC is a ∼21 kDa helical homodimeric lipoprotein expressed by *B. burgdorferi* during tick transmission and in the early stages of infection (16). OspC is implicated in facilitating spirochete egress from the tick during the course of a blood meal, enabling survival in the early stages of mammalian skin infection, and modulating transmigration across vascular walls (16-23). The highly immunogenic nature of OspC has lent itself to applications in Lyme disease serology-based diagnostics, including widely used commercial tests (24, 25). For example, the peptide derived from the conserved C-terminal 10 amino acids of OspC (peptide “C10”) is a component of the FDA approved diagnostic test *Borrelia* VlsE1/pepC10 from Zeus Scientific (Branchburg, NJ) (26-30). In addition to their diagnostic utility, linear B cell epitopes from different OspC types are a component of a widely used canine Lyme disease vaccine and being considered for human use (see references within (31)).

While multiple linear human B cell epitopes on OspC have been reported over the past three decades, differences in serologic assays, detection methodologies, and sample sizes from varying Lyme disease cohorts, makes it difficult to draw conclusions about relative reactivities of one peptide over another. This is problematic, because determining the relationships between OspC-derived peptides (including C10) and total OspC antibodies has practical implications for interpreting serologic assays. Moreover, there are >26 allelic variants or types of OspC reported in the United States, with amino acid sequence identities that range from 60-90% (32, 33). As different OspC types are associated with varying degrees of virulence and invasiveness, tracking epitope-specific responses to those particular types is of paramount importance (34, 35). In New York state, for example, OspC types A, B and K, which are associated with more invasive disease, represent ∼70% of all isolates (35-37).

With that in mind, we sought to validate human antibody reactivities to reported linear epitopes associated with OspC types A, B, and K to enable broad comparability of epitope usage across *B. burgdorferi* for diagnostic and vaccine development purposes.

## METHODS

### Chemicals and biological reagents

Chemicals and reagents were obtained from Thermo Fisher Scientific (Waltham, MA), unless noted otherwise. Buffers were prepared by the Wadsworth Center’s Cell and Tissue Culture core facility.

### Recombinant OspC and OspC-derived peptides

Recombinant *B. burgdorferi* OspC_A_ (residues 38 to 201; PDB ID 1GGQ; UniProt ID Q07337) (38), OspC_B_ (residues 38 to 202; *B. burgdorferi* strain ZS7; PDB ID 7UJ2) (39) and OspC_K_ (residues 38 to 202; *B. burgdorferi* strain 297; PDB ID 7UJ6) (40) were expressed in *E. coli* strain BL21 (DE3) and purified by nickel-affinity and size-exclusion chromatography, as described (41). Recombinant OspC_A_ with the C10 sequence (residues 201-210; PVVAESPKKP; “OspC_A_+C10”) was expressed and purified as above. Linear epitope prediction was done using Discotope 2.0 (42). OspC peptides (>80% purity) as described in **Table 1** were synthesized by Genemed Synthesis (San Antonio, TX) with a C-terminal GGGSK-biotin extension. We also synthesized the VlsE-derived C6 B31-17 peptide [MKKDDQIAAAIALRGMA] with the GGGSK-biotin linker (43).

**Table 1.**
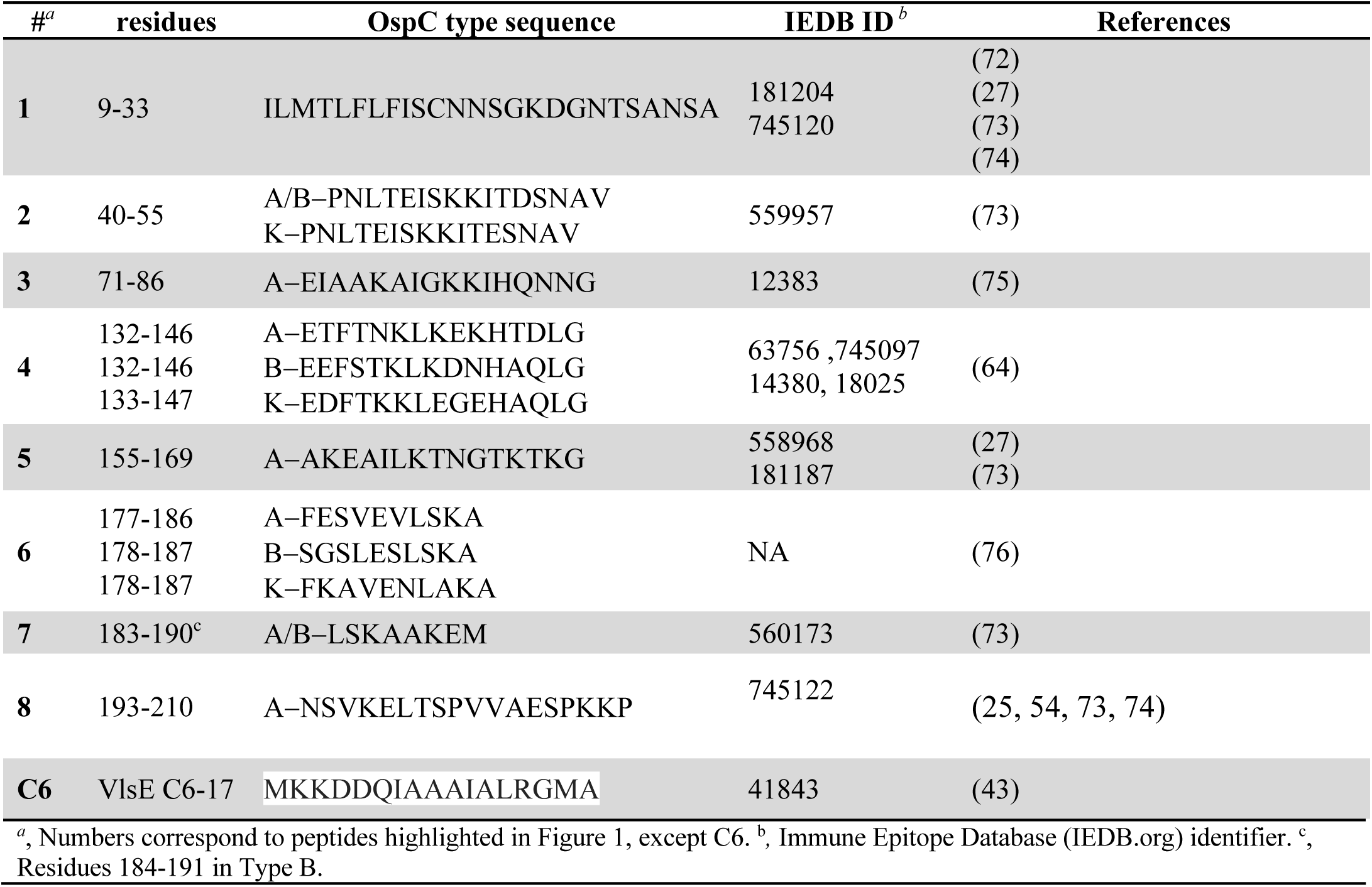
OspC and VlsE-derived peptides used in this study.

### Commercial and clinical LD serum samples

Commercial Lyme disease seronegative (Lot 10500586) and seropositive (Lot 10510438) pooled samples were used strictly as intra-assay and bead coupling confirmation controls throughout this study (ACCURUN products 810 and 130, respectively; SeraCare, Milford, MA). The Lyme disease seronegative samples (referred to as “controls” throughout the manuscript) consist of a commercial panel of 81 serum samples collected in 2017-2018 (Access Biologicals, Vista, CA). Five of the 81 serum samples were classified as extreme outliers for VlsE C6-17 reactivity by interquartile range (IQR) calculated using Microsoft Excel and therefore removed from the control panel. In the end, a total of 76 control samples were used as comparators with the diagnostic sample set and 75 samples were used as comparators for the PTLD sample set as a result of sample availability.

*B. burgdorferi* seropositive serum samples (n = 696) were obtained from the New York State Department of Health Wadsworth Center’s Diagnostic Immunology Laboratory. The samples had been submitted for Lyme disease serology and subjected to two-tiered testing consisting of (Tier 1) a C6 peptide screen (Immunetics®; C6 Lyme ELISA™) or Enzyme Linked Fluorescent Assay (ELFA; BioMerieux, VIDAS^®^ Lyme IgG II and Lyme IgM II; Durham, NC), followed by (Tier 2) IgM and IgG detection by Western blot (MarDX^®^; Trinity Biotech, Carlsbad, CA). *B. burgdorferi*-specific IgM reactivity was defined as ≥2 positive bands, with IgG reactivity defined as ≥5 positive bands. Serum samples were aliquoted, de-identified, and classified as IgM-positive/IgG-negative, IgM-positive/IgG-positive, or IgM-negative/IgG-positive, based on the Western blot results. Post-treatment Lyme disease (PTLD) serum samples (n=158) were kindly provided by the Lyme Disease Biobank at Nuvance Health^®^ (Danbury, CT). PTLD was defined as described by Aucott (8).

### Multiplexed OspC and OspC peptide microsphere immunoassays (MIA)

Recombinant OspC_A_, OspC_B_, or OspC_K_ (5 μg) were coupled to Magplex-C microspheres (1 x 10^6^) using sulfo-NHS (N-hydroxysulfosuccinimide) and EDC [1-ethyl-3-(3-dimethylaminopropyl) carbodiimide hydrochloride], as recommended by the manufacture (Luminex Corp., Austin, TX). Coupled beads were diluted in xMap^®^ AbC Wash Buffer to a concentration of 5 x 10^6^ beads/ml. Biotin-labeled peptides were complexed to Magplex^®^-avidin microspheres following protocols provided by the manufacturer (Luminex Corp.). Microspheres were resuspended in 250 μL of assay buffer (1 x PBS, 2% BSA, pH 7.4) then subjected to vortexing and sonication. A total of 1 x 10^6^ beads in assay buffer was mixed with biotin-conjugated peptides (∼5 μg) and incubated for 30 min at room temperature. The microsphere suspensions were then washed three times using wash buffer (1 x PBS, 2% BSA, 0.02% TWEEN-20, 0.05% Sodium azide, pH 7.4) and a magnetic separator, resuspended in 500 μL of assay buffer, and stored at 4°C until use. Successful coupling of OspC and peptides to the Magplex-C microspheres was confirmed by reactivity with immune serum and/or monoclonal antibodies.

For experimental use, assay buffer was used to dilute bead stocks (1:50) and human serum (1:100). The bead dilution (50 μL) and diluted serum (50 μL) were combined in black, clear-bottomed, non-binding, chimney 96-well plates (Greiner Bio-One, Monroe, North Carolina) and allowed to incubate for 60 min in a tabletop shaker (600 rpm) at room temperature. Plates were washed three times using a magnetic separator and wash buffer. Secondary antibody goat anti-Human IgG Fc, eBioscience (Invitrogen, Carlsbad, CA) was diluted 1:500 in assay buffer and added (100 μL) to each well. The secondary antibody was allowed to incubate for 30 min in a tabletop shaker (600 rpm) at room temperature. Plates were washed as stated above and beads were resuspended in 100 μL of wash buffer. Samples were analyzed using a FlexMap 3D (Luminex Corp). We defined reactivity as the mean MFI +6 SD, based on previous studies conducted in the Wadsworth Center’s clinical laboratories (44, 45). Index values were calculated using the sample MFI divided by the reactive cutoff for each bead set (*i.e.*, each peptide coated bead versus itself) such that an index values of >1.0 indicates positive reactivity for a given bead set.

### Statistical analysis

Statistical analyses were performed using R 4.3.0 with R packages readxl and tidyverse (46-48). MFIs (log_10_) were first subjected to the Shapiro-Wilk’s test to assess normality, then either Levene’s or Fligner-Killeen tests to compare variances. Mann-Whitney U tests with an alpha level of 0.05 were used to determine statistical significance for reactivity determination of OspC subtypes and OspC-derived peptides. R package corrplot (49) was used to generate the Spearman’s Rank correlation matrix. R package pheatmap was used to create the hierarchically clustered heatmaps (50). R packages ggthemes [https://github.com/jrnold/ggthemes], RColorBrewer [https://cran.r-project.org/web/packages/RcolorBrewer/index.html], and ggpubr [https://CRAN.R-project.org/package=ggpubr] were used in formatting.

### Molecular modeling

PyMol (PyMOL Molecular Graphics System, Version 3.0 Schrödinger, LLC) was used for epitope modeling using OspC PDB ID: **1GGQ** (strain B31, OspC_A_).

## RESULTS

We sought to validate the reactivity of OspC linear B cell epitopes, including those already accessioned in the Immune Epitope Database (iedb.org), that have been directly or indirectly implicated in Lyme disease diagnostics and/or immunity to *B. burgdorferi*. Eight different epitopes, including the C10 peptide, were chosen for analysis **(Table 1)**. When localized onto the structure of OspC_A_ using PyMol, the epitopes represent ∼30% of the surface area of the molecule (**Figure 1A**). An alignment of the primary amino acid sequences of the three OspC types (A, B, K) associated with invasive disease in northeast United States revealed polymorphisms in a number of these epitopes (**Table 1**; **Figure 1B**) (51, 52). For this reason, a total of 13 different peptides were generated to encompass the specific amino acid sequences for each epitope within OspC_A_, OspC_B_, and OspC_K_ (**Table 1**). The peptides were synthesized with a C-terminal biotin tag and coupled to streptavidin-coated microspheres for multiplex immunoassays (MIA) by Luminex. As a positive control, we also coupled microspheres with the VlsE C6 B31-17 peptide from *B. burgdorferi* strain B31(43).

**Figure 1.**
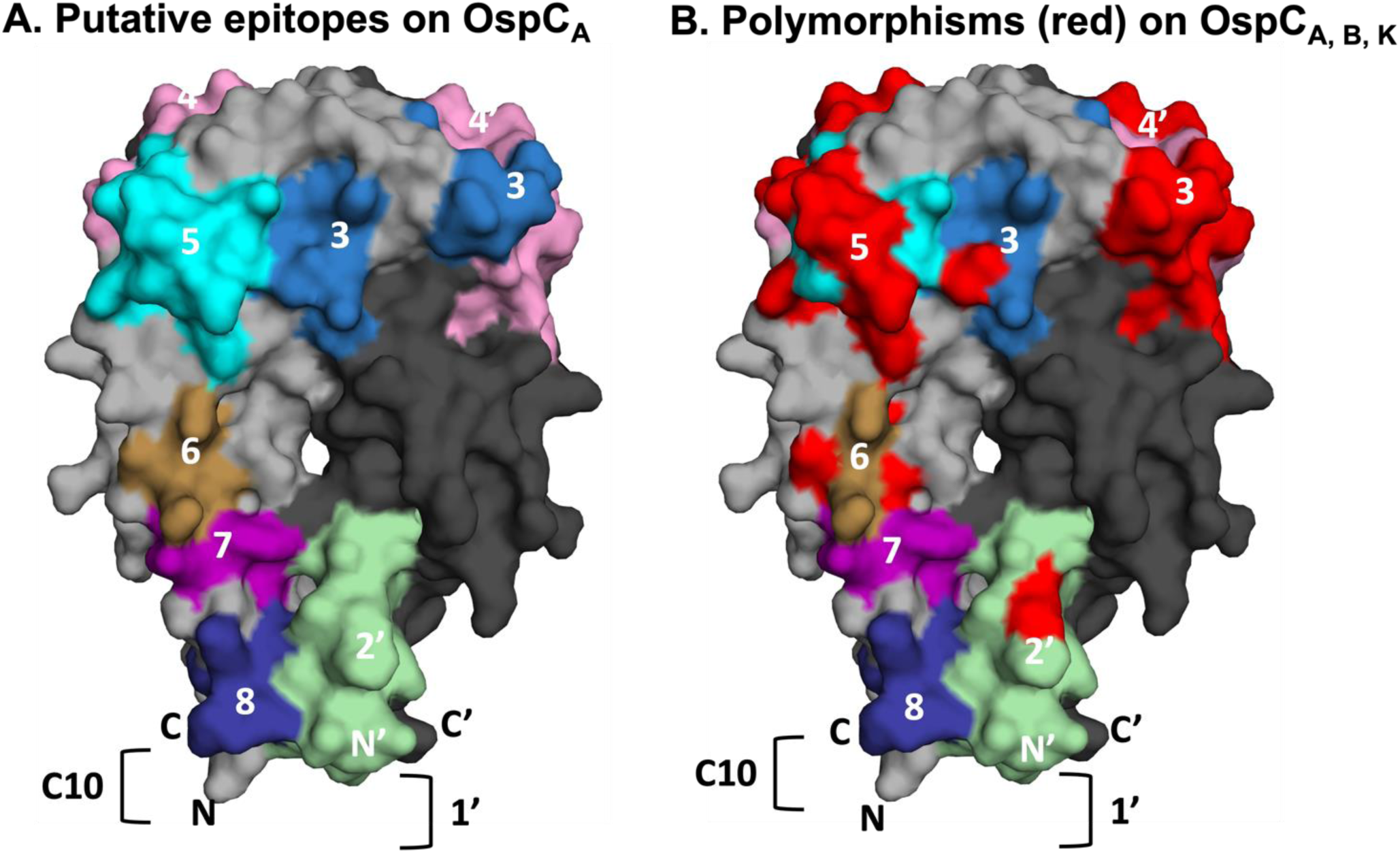
Relative location of linear epitopes on homodimeric OspC_A_. (**A**) Surface representation of homodimeric OspC_A_ (residues 38-201; PDB 1GGQ] with one monomer (OspC) colored gray and the other, denoted with an apostrophe (OspC’), colored in charcoal. The linear epitopes examined in this study are coded and numbered according to **Table 1**. Epitope 1 is represented as a bracket (]) on OspC’, as its corresponding residues were truncated in the recombinant version of OspC_A_ used for X-ray crystal structure analysis. Similarly, only residues 193-201 of epitope 8 (residues 193-210) are shown because the C-terminus of OspC_A_ was truncated at position 201. Overlap between epitopes 6 and 7 are colored as epitope 6. (**B**) The same image as Panel A except that amino acid differences between OspC types A, B and K within the eight epitopes are colored red to illustrate degree of polymorphisms.

We employed two *B. burgdorferi* seropositive serum sample collections to gauge the relative reactivity of recombinant OspC, OspC-derived peptides, and the VlsE C6-17 peptide. A “diagnostic” collection consisted of 696 de-identified human serum samples from the Wadsworth Center’s Diagnostic Immunology laboratory classified as *B. burgdorferi* seropositive and categorized based on Western blot banding profiles, as IgM^+^/IgG^-^ (+/-), IgM^+^/IgG^+^ (+/+), or IgM^-^ /IgG^+^ (-/+). A second collection provided by the Lyme Disease Biobank at Nuvance Health consisted of 158 serum samples from PTLD patients. These two collections were compared to a commercial panel consisting of ∼75 serum samples confirmed negative for VlsE C6-17 peptide reactivity. We examined both serum IgG and IgM (see Supplemental Information) reactivity profiles.

We found that the diagnostic immunology samples were significantly more reactive with recombinant OspC than the control samples, although reactivity was skewed towards OspC_A_ and OspC_B_ more than OspC_K_ **(Figure 2A**). In terms of peptides, the diagnostic serum samples were on average significantly more reactive than control serums samples with all the OspC-derived peptides tested, except for peptides 71-86A and 178-187B (**Figure 2A**). The average serum IgG reactivity was highest against 193-210_A_ (“C10”) and VlsE C6-17 (p<0.0001) with the remainder of the peptides being low to moderately reactive relative to the control. Analysis of IgM antibodies revealed significant reactivity with OspC types A, B and K but considerably less reactivity with OspC-derive peptides. Significant IgM reactivity in the diagnostics samples relative to controls was only observed for peptides 9-33_ABK_, 155-169_A_, 178-187_B, K_, and 193-210_A_ (“C10”) (**Figure S1**).

**Figure 2.**
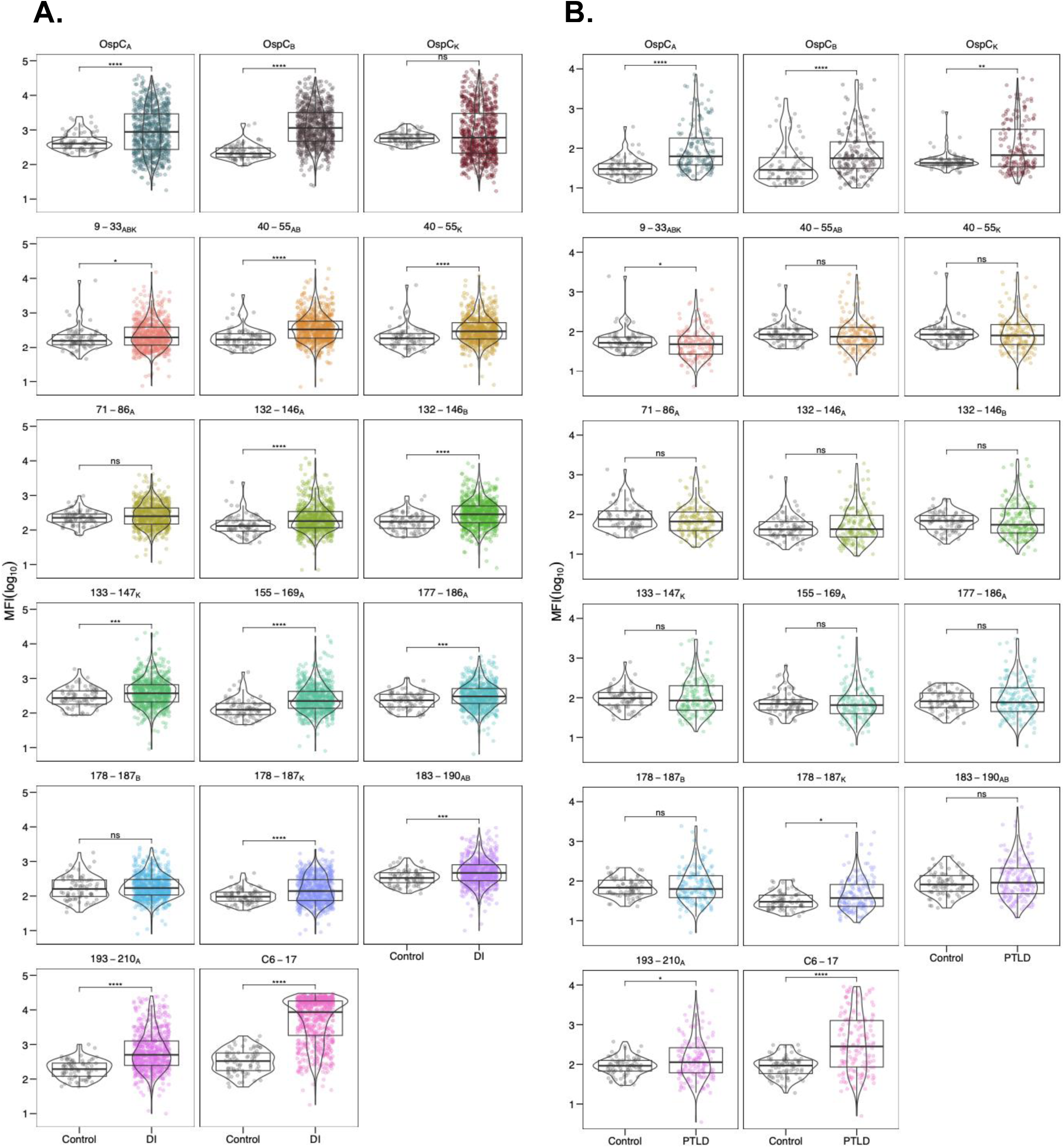
Serum IgG reactivity of diagnostic and PTLD samples with OspC and OspC-derived peptides. (**A**) DI and (**B**) PTLD serum samples (diluted 1:100) were subject to MIA using microspheres coated with recombinant dimeric OspC types A, B and K (top rows), OspC-peptides as described in Table 1, and C6-17 peptide. Panels are labeled by corresponding residue numbers for each OspC type. MFI values were log_10_ transformed and compared to the control sample set. Significance was determined by the Mann-Whitney U-test (*, p< 0.05).

To apply a more rigorous evaluation of peptide reactivity, MFI values were subjected to a cutoff value of ≥6 SD above the mean of the control MFI, log_10_ transformed, then plotted to reveal both the number of serum samples above the cutoff (% positive) as well as the magnitude of samples over the cutoff (fold increase). We also derived box and whisker plots to illustrate the upper and lower quartiles as well as the median reactivity to each antigen/peptide with superimposed violin plots to outline sample distribution. Using these metrics, reactivity with OspC_A_, OspC_B_ and OspC_K_ ranged from 24-42% of the diagnostic samples with the fold increase ranging from 2.8 to 4.7 (**Table 2**; **Figure 3A**). In terms of peptides, VlsE C6-17 was the most reactive (71% positive; fold increase 5.8), followed by 193-210_A_ (“C10”) (26% positive; fold increase 4.4) (**Table 2**; **Figure 3B**). The percent of samples >6SD for the other 12 peptides ranged from 0.1 to 10% with the fold increase between 1.2 and 2.5 (**Table 2**; **Figure 3B**). Lowering the cutoff threshold to >3 SD resulted in higher overall peptide reactivities, as expected, but the pattern remained the same as >6SD (**Table S1**). Thus, within a given cohort of *B. burgdorferi* seropositive samples, serum IgG reactivity to OspC-derived peptides (apart from 193-210_A_) is limited to a small subset of individuals, which has important consequences for use of such peptide in diagnostic applications. In terms of IgM reactivity, the trends were similar to IgG except that the most pronounced antibody interactions were with peptides 177-186_B, K_ followed by 155-169_A_ and 193-201_A_ (**Table S2**). The differential peptide reactivity profiles between IgG and IgM might have diagnostic utility.

**Figure 3.**
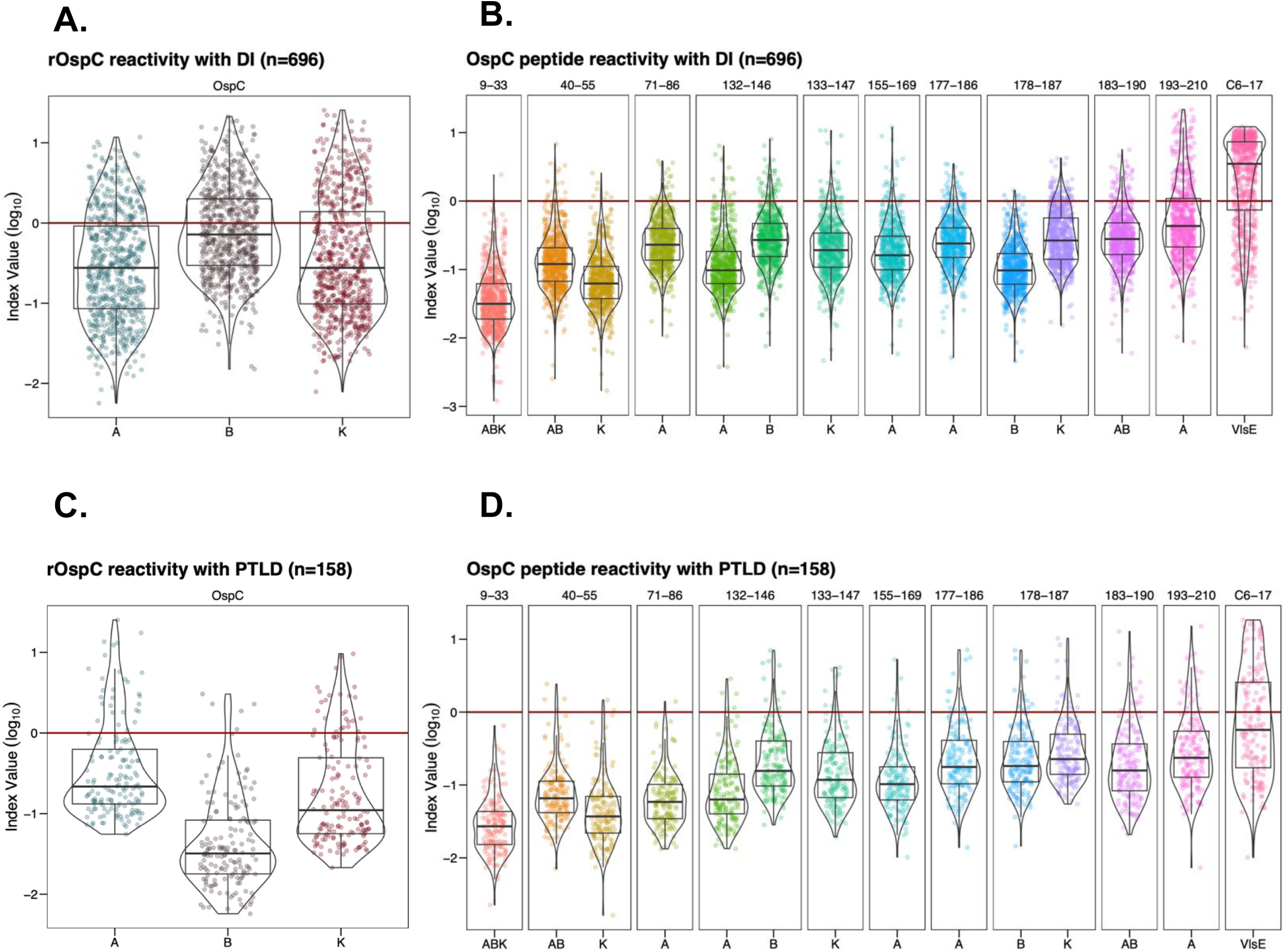
Indexed (>6SD) IgG reactivity of DI and PTLD serum samples with OspC and OspC-derived peptides. MFI values for OspC and OspC- and VlsE-derived peptides were indexed using a cutoff of 6SD above mean control reactivity. The resulting values were then log_10_ transformed with each point representing a single sample. OspC subtypes are listed along the x-axis, residues are indicated along the top of the plot (**B**, **D**), and each antigen is represented by a different color. In each of the graphs, the 6 SD cutoff corresponds to a value of 0 on the y-axis and is depicted with horizontal red line. All samples falling above y = 0 were found to be positive for the given antigen and were used to calculate percent positivity and fold increase as reported in Tables 2 and 3. Box and whisker plots illustrate the upper and lower quartiles as well as the median reactivity to each antigen (thick center line). Sample points that fall above or below the whiskers are considered outliers. Violin plots outline the distribution of data.

**Table 2:** Serum IgG reactivity > 6SD to OspCA/B/K and OspC peptides in diagnostic and PTLD serum samples.

| OspC or peptide | Diagnostic |  | PTLD |  |
| --- | --- | --- | --- | --- |
|  | n = (%)<sup>a</sup> | Fold increase (SD)<sup>b</sup> | n = (%)<sup>a</sup> | Fold increase (SD)<sup>b</sup> |
| OspC<sub>A</sub> | 167 (24.0) | 2.8 (1.9) | 29 (18.4) | 5.5 (5.6) |
| OspC<sub>B</sub> | 287 (41.2) | 3.7 (3.4) | 5 (3.2) | 2.1 (0.6) |
| OspC<sub>K</sub> | 195 (28.0) | 4.7 (4.3) | 26 (16.5) | 2.8 (2.2) |
| 9-33<sub>ABK</sub> | 1 (0.1) | 2.4 (N/A) | 0 (0) | N/A (N/A) |
| 40-55<sub>AB</sub> | 17 (2.4) | 2.1 (1.4) | 4 (2.5) | 1.6 (0.5) |
| 40-55<sub>K</sub> | 7 (1.0) | 1.5 (0.5) | 2 (1.3) | 1.3 (0.2) |
| 71-86<sub>A</sub> | 28 (4.0) | 1.7 (0.8) | 1 (0.6) | 1.4 (NA) |
| 132-146<sub>A</sub> | 18 (2.6) | 2.4 (1.5) | 5 (3.2) | 1.7 (0.6) |
| 132-146<sub>B</sub> | 48 (6.9) | 2.0 (1.3) | 11 (7.0) | 3.0 (1.9) |
| 133-147<sub>K</sub> | 36 (5.2) | 2.4 (2.6) | 8 (5.1) | 2.5 (1.2) |
| 155-169<sub>A</sub> | 30 (4.3) | 2.5 (2.5) | 5 (3.2) | 2.5 (1.5) |
| 177-186<sub>A</sub> | 38 (5.5) | 1.6 (0.7) | 13 (8.2) | 2.6 (2.0) |
| 178-187<sub>K</sub> | 70 (10.1) | 1.5 (0.7) | 15 (9.5) | 3.0 (2.8) |
| 178-187<sub>B</sub> | 7 (1.0) | 1.2 (0.1) | 12 (7.6) | 2.7 (1.9) |
| 183-190<sub>AB</sub><sup>c</sup> | 53 (7.6) | 1.9 (1.0) | 14 (8.9) | 3.0 (3.1) |
| 193-210<sub>A</sub> (C10) | 185 (26.6) | 4.4 (4.5) | 22 (14.0) | 3.3 (3.0) |
| VlsE C6-17 | 496 (71.3) | 5.8 (2.9) | 64 (40.5) | 5.6 (5.0) |

<sup>a</sup>, The number of samples (n =) and percent reactivity (“%”) to each antigen in the Diagnostic (n=696) and PTLTD (n=158) sample sets were calculated using a cutoff of >6SD above the mean of controls samples; <sup>b</sup>, The fold increase (magnitude) of positive sample antibody reactivity over controls such than an index value of 1 is 6SD above the control mean, while 2.8 corresponds to ~16SD greater than the mean control. <sup>c</sup>, OspC<sub>B</sub> residues 184-191.

The same overall serologic analysis was performed with a panel of PTLD samples. The PTLD samples were significantly more reactive with OspC_A_, OspC_B_ and OspC_K_ than the control serum samples (**Figure 2B**). For OspC_A_ and OspC_K_, 16 and 18% of the samples were above the ≥6SD cutoff with the fold increase between 2.8-5 (**Table 2**; **Figure 3C**). In the case of OspC_B_, only 3.2% were above the cutoff with a fold increase of 2.1 (**Table 2**; **Figure 3C**). In terms of linear epitopes, reactivity significantly above controls was limited to VlsE C6-17 (40% positive; fold increase 5.6), 193-210_A_ (“C10”) (14% positive; fold increase 3.3), and 178-187_K_ (9.5%; positive; fold increase 3.0) (**Table 2**: **Figure 2B**; **Figure 3D**). Comparatively, PTLD samples had significantly less reactivity to peptide 9-33_ABK_ relative to controls (**Figure 2B**). Only a small fraction of the PTLD serum samples were found to be positive for the remaining 10 peptides (range 1.3-9.5%) and the MFIs did not achieve statistical significance (**Table 2**; **Figure 2B**). PTLD samples were plotted to visualize OspC and peptide reactivities relative to the cutoff (**Figure 3C-D**). There is notably less reactivity to both 193-210_A_ (“C10”) and C6-17 in the PTLD cohort in comparison to the diagnostic population (**Figure 3B, 3D**).

Understanding the relationships between OspC and OspC-derived peptide epitopes has implications for Lyme disease diagnostics and possibly immunity to *B. burgdorferi*. We expected that antibody reactivity with the OspC-derived peptides would strongly correlate with overall OspC antibody levels in any given sample, suggesting that the two populations track with each other and serve as redundant indicators of a previous *B. burgdorferi* infection. To investigate these relationships, we generated correlation matrices (Spearman’s Rank) showing the correlation between OspC and the each of the different OspC-derived peptides, including C10 and C6-17 (**Figure 4**). While all correlations were determined to be positive, in the diagnostic samples, C10 reactivity was only weakly correlated with OspC_A_, OspC_B_, or OspC_K_ [r_s_ = 0.2-0.4] (**Figure 4A**). Rather, C10 reactivity was moderately to strongly correlated with other OspC-derived peptides, including 132-146_A_ (r_s_ = 0.63), 183-190_AB_ (r_s_ = 0.58), and 40-55_ABK_ (r_s_ = 0.58). Indeed, inter-peptide correlations were generally strong across the board (e.g., 132-146_B_ versus 177-186_A_, r_s_ >0.9). A similar pattern was observed in the PTLD samples: OspC-derived peptide reactivities were strongly correlated with each other (r_s_ = 0.63-0.98), but weakly correlated with OspC_A_, OspC_B_, and OspC_K_ (r_s_ = 0.2-0.4) (**Figure 4B**). In the case of VlsE C6-17, reactivity was weakly correlated with OspC and OspC-derived peptides in both the diagnostic and PTLD samples, except for moderate correlativity with 193-210_A_ (“C10”) in the PTLD cohort (r_s_ = 0.56). These results reveal a significant degree of discordance (asynchrony) among circulating antibody levels against OspC, OspC-derived peptides and the VlsE C6-17 peptide in both diagnostic and PTLD cohorts.

**Figure 4.**
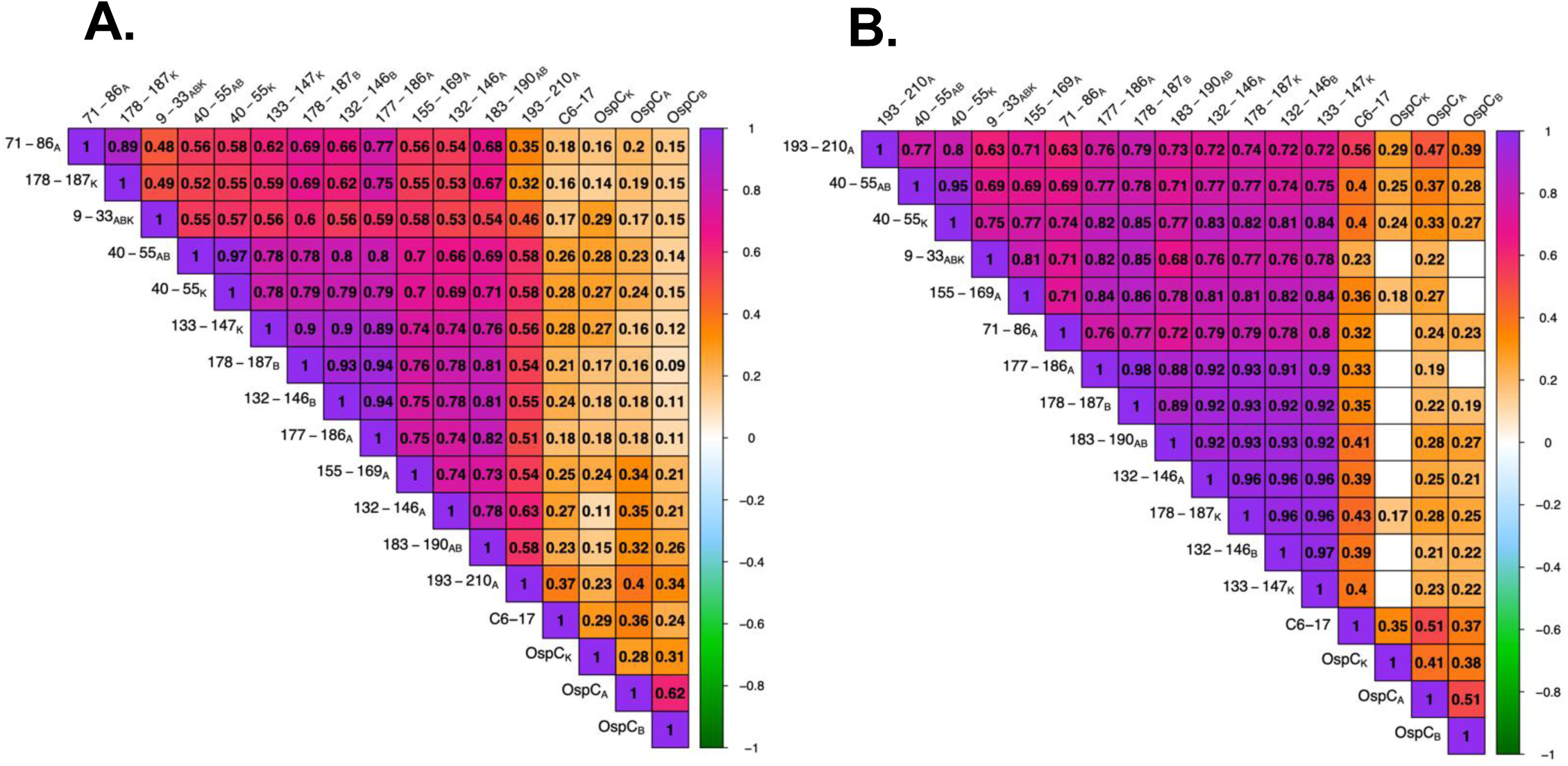
Correlation matrices of rOspC and OspC-derived peptide reactivity in diagnostic and PTLD serum. Diagnostic (**A**) and PTLD (**B**) serum sample MFIs (indexed and log_10_ transformed) were subjected to Spearman’s Rank correlation. Only significant correlation coefficients (r_s_) are displayed (confidence level = 0.95). The degree and direction of correlation is indicated by the color scale. Interpretations of the strength of each correlation was conducted as such: >0.7 indicates a strong correlation, 0.5 – 0.7 is moderate, 0.3 – 0.5 is weak, and 0.0 – 0.3 is negligible.

The absence of a strong correlation between antibody levels to OspC and OspC-derived peptides led us to subject the diagnostic and PTLD datasets to hierarchical clustering (HC) as means to identify possible relationships in immunoreactivity profiles. In the case of the diagnostic samples, the horizontal (top) dendrogram revealed three distinct branches consisting of (1) OspC_A_, OspC_B_, and OspC_K_, (2) C6-17 and 193-210_A_ (“C10”), and (3) the other OspC-derived peptides (**Figure 5A**). In cluster 1 (OspC_A_, OspC_B_, OspC_K_), high MFI values were distributed throughout the column with a concentration towards the lower quartile. There was no obvious clustering on the vertical dendrogram to indicate a relationship between OspC reactivity and seropositivity categories (i.e., IgM^+/-^, IgG^+/-^).

**Figure 5.**
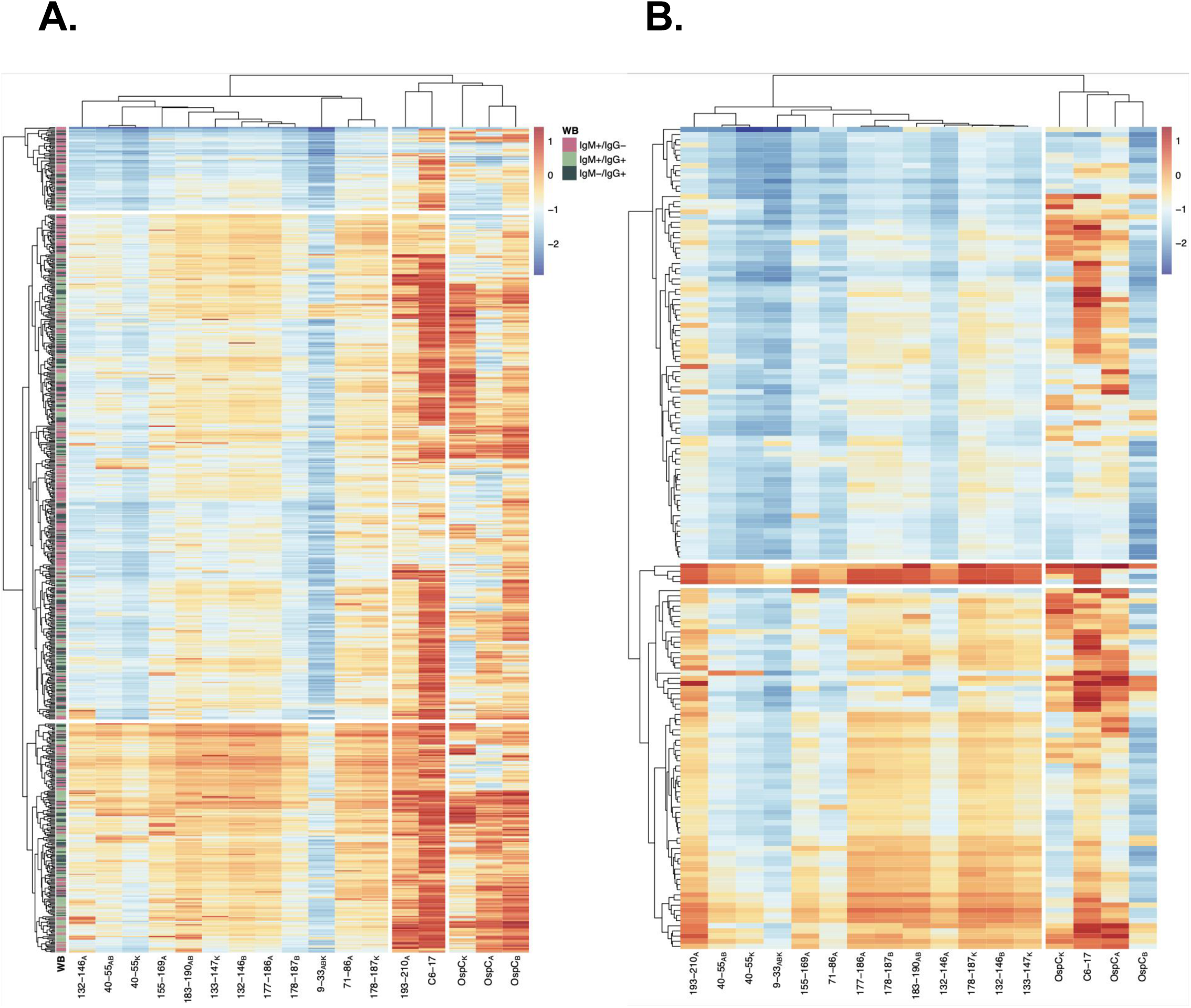
Hierarchical clustering of rOspC and OspC-derived reactivity in diagnostic and PTLD serum. Heatmaps were generated using diagnostic (**A**) and PTLD (**B**) serum sample MFIs (indexed and log_10_ transformed). Rows represent individual samples and columns represent the panel of antigens. The color scales of **A** and B are identical to one another (allowing for direct comparison), with 0 representing the positivity cutoff (as seen in Figure 3). Annotation of the diagnostic samples (**A**) with Western blot profile data (previously described in the Materials and Methods) is designated by the legend. The columns were subjected to hierarchical clustering by correlation and the rows were clustered by Euclidean distance to better visualize patterns in reactivity profiles.

In branch 2, high VlsE C6-17 reactivity was present throughout the column, albeit, with three hot (orange and red shading) zones and one cold (deep blue shading) zone. In the 193-210_A_ (“C10”) column, there was generally an even distribution of MFI intensities only occasionally punctuated with hot spots that often mirror 193-210_A_ (“C10”) reactivity. Samples that were “cold” for C6-17 and 193-210_A_ (“C10”) tended to be within the IgM+/IgG-pool, possibly indicative of an early response to *B. burgdorferi* infection.

There were several interesting patterns within branch 3, which consisted of OspC-derived peptides: uniformly <u>low reactivity</u> with peptide 9-33_ABK_, as noted in Figures 2 and 3; a subpopulation of samples that were uniformly <u>nonreactive</u> with all the OspC peptides, as evidenced by a blue band across the top of the figure, which in some instances spilled over to 193-210_A_ (“C10”) and VlsE C6-17 columns; and, finally, a subpopulation of samples with <u>moderate to high</u> pan-OspC peptide reactivity, as evidenced by a horizontal band of orange/red shading situated in the lower third of the figure. Collectively, the HC analysis reveals a remarkable degree of discordance in serum antibody profiles associated with *B. burgdorferi* seropositivity within the diagnostic samples.

The HC plot of the PTLD serum samples was clearly distinct from the diagnostic samples (**Figure 5B**). On the horizontal dendrogram, for example, there were just two major branches not three: one encompassing OspC_ABK_ and C6-17, and the other containing the OspC-derived peptides, including 193-210_A_ (“C10”). On the vertical dendrogram, there were two branches: the top branch corresponding to low peptide reactivity with or without concomitant C6-17 and OspC reactivities; and the bottom branch corresponding to high peptide reactivity that were more frequently than not associated with high C6-17 reactivity. Within the bottom branch, there was a notable subbranch corresponding to 4 samples with strikingly high reactivity with C10, C6-17, and most of the OspC-derived peptides (including 9-33_ABK_). We would tentatively classify these handful of samples as “polyreactive” (53).

### Contribution of C10 to overall OspC antibody reactivity

There is some debate as to what proportion C10-specific antibodies constitute relative to total OspC antibodies, with some studies reporting that C10-specific antibodies make up an overwhelming proportion (54), while others suggesting the opposite (55). As our prior analysis included truncated derivatives of OspC that lacked C10, we expressed a second version of OspC in which residues 202-210 (VVAESPKKP) were included. We then revisited Luminex analysis in which we compared MFI to OspC_A_ versus OspC_A_+C10. MFIs for OspC_A_+C10 were significantly greater than OspC_A_ in both diagnostic and PTLD cohorts, demonstrating that antibody reactivity to the conserved C-terminal residues constitute an important component of the overall antibody response (p <0.05; Mann-Whitney U test) (**Figure 6A, 6D**). We then investigated the relationship between C10 reactivity and OspC_A_ vs OspC_A_+C10 using a linear regression model for diagnostic samples (**Figure 6B, 6C**) and PTLD samples (**Figure 6E, 6F**). There is a slight increase (0.0702) in the adjusted R-squared value between the C10 peptide MFI and OspC_A_+C10, as compared to OspC_A_ for the DI cohort (**Figure 6B, 6C**). A similar increase (0.0848) in the adjusted R-squared value was observed between the C10 peptide and OspC_A_+C10 in comparison to OspC_A_ for the PTLD population. However, the overall adjusted R-squared value was negligible, thereby demonstrating that C10-specific antibodies constitute only a small fraction of the total OspC reactivity.

**Figure 6:**
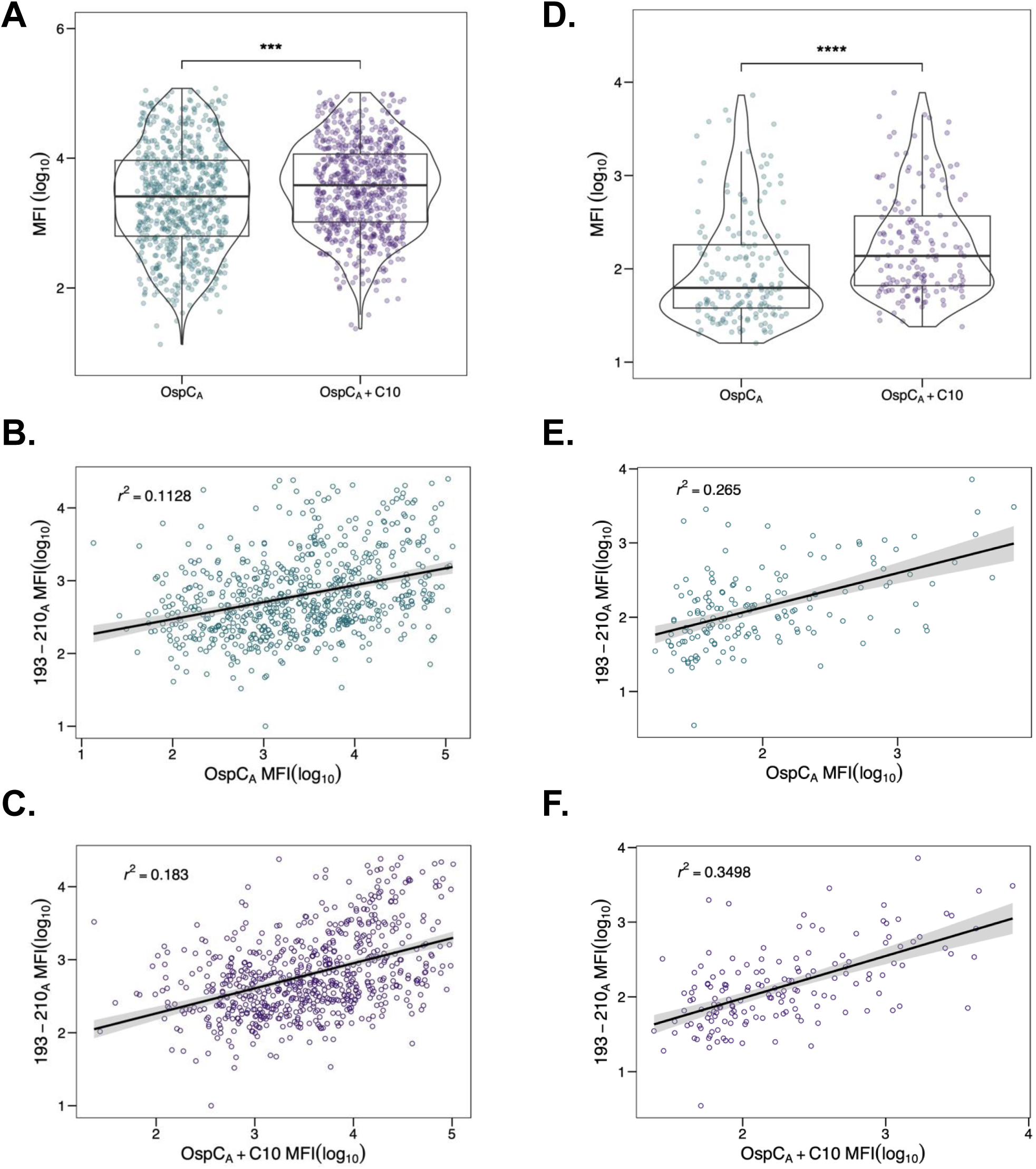
Correlation between OspC_A_ reactivity and C10 region. Diagnostic (n=686) and PTLD (n=158) samples were subjected to Luminex with OspC_A_, OspC_A_+C10, and 193-210_A_ as described in the Materials and Methods. OspC_A_ MFIs were evaluated against OspC_A_+C10 for diagnostic **(A)** and PTLD **(D)** samples. Significance was determined by Mann-Whitney U test. *, p < 0.05. To examine the contribution of the C10 region to OspC_A_ reactivity, MFIs for peptide 193-210_A_ were plotted (on the y-axis) versus MFIs from OspC_A_ and OspC_A_+C10 (on the x-axis) for diagnostic samples **(B, C)** and PTLD samples **(E, F)**. Best fit lines and confidence intervals (level = 0.95) were added to each plot using a linear regression model. Adjusted R-squared (*r^2^*) values were calculated in R.

## DISCUSSION

Defining the breadth and the frequency of linear B cell epitope reactivity on OspC has important implications for both Lyme disease diagnostics and vaccines. In this report, a panel of peptides representing eight previously reported and predicted linear epitopes on OspC types A, B and K, were examined by multiplex immunoassay for serum IgG reactivity in diagnostic (n∼700) and PTLD (n∼150) *B. burgdorferi* seropositive sample sets. The results revealed that seven of the eight peptides are indeed antibody targets even though reactivity to any given peptide may be confined to a fraction (2-10%) of the total samples tested. The exception being peptide C10, which was 26% positive (4.4 positive fold increase value) in the diagnostic sample set and 14% positive (3.3 positive fold increase value) in the PLTDs sample set. Interestingly, in the diagnostic sample set, C10 antibody titers did not correlate with either reactivity to OspC itself or any of the other OspC-derived peptides. In the PTLD sample set, C10 antibody levels did correlate with the other OspC-derived peptides but were distinct from OspC. These results have clinical implications, as they demonstrate that the human B cell response to linear epitopes on OspC is both variable and asynchronous.

The C10 peptide corresponds to the stretch of ∼10 highly conserved amino acids at C-terminus of OspC, first recognized as an immunodominant epitope in neuroborreliosis patients (54). Since then, the C10 peptide and derivatives have been integrated into numerous diagnostic assays, including the Zeus VlseE1/C10 ELISA (25, 27, 28). Dolange and colleagues recently proposed the use of high affinity C10-specific monoclonal antibodies as tools to detect circulating levels of *B. burgdorferi* antigens (56). While there is little debate as to whether C10 constitutes a human B cell epitope, the issue of whether antibodies to C10 contribute to borreliacidal activity remains contentious. Lovrich and colleagues reported that OspC-specific borreliacidal activity (in a handful of human serum samples) was almost entirely directed against C10 (57, 58), whereas Izac and colleagues argued that, in rodents, canines and non-human primates, C10-specific antibodies contribute little if any borreliacidal activity (59). Although our results do not resolve this issue per se, they do raise questions about the underlying B cell biology associated with antibodies to C10 and OspC. The fact that C10 and OspC antibody titers did not correlate with each other in the two serum sample sets examined suggests differences in the kinetics of antibody induction, circulating half-life, and/or plasma cell persistence. Others have suggested, at least in the case of VlsE, there is a distinct progression of epitope reactivity over the course of *B. burgdorferi* infection with readily accessible epitopes targeted first and more obstructed, membrane proximal epitopes following (60). Resolving detailed questions about the nature of the human B cell response to OspC is becoming more feasible as single cell V_H_ and V_L_ repertoires are catalogued over the course of Lyme disease (11, 61).

B cell epitope prediction software like Discotope 3.0 identifies residues 133-147 as having a high immunogenic propensity (62). That prediction was borne out in our study, as evidenced by statistically significant reactivity of serum samples with the three peptides corresponding to residues 132-146_AB_ and 133-147_K_. Others have reported similar findings in mice and in a few human samples. Namely, Earnhart recognized residues 136-150 (which they refer to as “loop 5”) as a target of antibodies in *B. burgdorferi*-infected mice (36). In a follow-up study, the same investigators identified two serum samples from early Lyme disease patients that reacted with peptides spanning 136-144 (63). A subsequent report by Arnaboldi and colleagues did not detect binding to three different 15-mer OspC_K_-derived peptides spanning residues 131-155, possibly because their sample size was limited (27). In our collection of almost 700 diagnostic samples, the frequency of peptide positivity was admittedly relatively low (3-7%). Nonetheless, antibodies targeting this epitopic region may be consequential in terms of promoting spirochete clearance (36). For example, the potent borreliacidal mouse monoclonal antibody 16.22 recognizes OspC_A_ residues 133-147 with the key contact points postulated to be residues 139-141 (“KXK” motif) (64). Thus, a more detailed examination of B cell epitopes within this region of OspC in human Lyme disease patients is warranted.

It has been postulated that serum antibody profiling at the antigen and possibly even epitope level may be able to resolve different stages of Lyme disease (e.g., acute, acute resolved, PTLD) (10, 60). While the antibody profiling in our study was limited to serum IgG reactivity to OspC, OspC-derived peptides, and VlsE C6-17, the results revealed distinct patterns between the diagnostic and PTLD sample sets when subjected to HC. Most notable was the classification of the C10 peptide: in the diagnostic sample set C10 clustered with C6-17, whereas in the PTLD samples it clustered with the other OspC-derived peptides. Thus, it is tempting to speculate that the ratio of C10 to C6-17 serum IgG reactivity may have some utility in defining disease progression. By the same token, as virtually all samples tested were positive for either C10 and/or C6-17, establishing a combined C10 + C6-17 threshold value might serve to alleviate false positives in Lyme disease diagnostics based solely on serology (65, 66). That said we recognize that any claims to this effect are premature, considering that the serum samples employed in our study are not associated with any clinical information beyond seropositivity. Moreover, we lack a “post-Lyme healthy” cohort that Chandra and colleagues so effectively used in their study to compare with their PTLD cohort (10).

One interesting grouping that emerged from the PTLD cohort was a cluster of four individuals who demonstrated pan-reactivity with the OspC-derived peptides and VlsE C6-17 peptide. In follow-up analysis, those same four samples in question also reacted with other *B. burgdorferi*-derived peptides, as well as control non-*B. burgdorferi*-derived peptides, but not necessarily with the corresponding recombinant proteins (G. Freeman-Gallant, unpublished results). Thus, the serum antibodies from those particular individuals seemingly have a propensity for peptide antigens, possibly due to a particular bias in germline chain usage and corresponding paratopes conducive to accommodating peptides (67). Whether or not there is a germline bias in antibodies associated with PTLD is worthy of investigation, especially in light of finding related to COVID-19. Moreover, the availability of bulk plasmablast BCR sequences, as well as single cell paired BCR sequences from LD patients opens the door to detailed antibody reactivity profiles and links to disease manifestations (68). We are also exploring the possibility that these particular samples may have profiles consistent with autoantibodies (69) or have characteristics associated with polyreactivity that might promote ligation to spirochete surface antigens (70, 71).

## Acknowledgements

We thank Dr. William Lee and other members of the Diagnostic Immunology Laboratory at the Wadsworth Center for providing residual diagnostic serum samples. We gratefully acknowledge Dr. John Martignetti and Lisa Arrigo of the Nuvance Health Lyme disease Biobank for generously providing PTLD serum samples. We thank Elizabeth Cavosie (Wadsworth Center) for administrative assistance and Greta Van Slyke (Wadsworth Center) for establishing Luminex protocols. This work was supported by the National Institute of Allergy and Infectious Diseases (NIAID), National Institutes of Health, Department of Health and Human Services, Contract No. 75N93019C00040 (PI/PD Mantis). The content is solely the responsibility of the authors and does not necessarily represent the official views of the National Institutes of Health.

## References Cited

1. Kugeler, K. J., A. M. Schwartz, M. J. Delorey, P. S. Mead, and A. F. Hinckley. 2021. Estimating the Frequency of Lyme Disease Diagnoses, United States, 2010-2018. Emerg Infect Dis 27: 616-619.

2. Bobe, J. R., B. L. Jutras, E. J. Horn, M. E. Embers, A. Bailey, R. L. Moritz, Y. Zhang, M. J. Soloski, R. S. Ostfeld, R. T. Marconi, J. Aucott, A. Ma’ayan, F. Keesing, K. Lewis, C. Ben Mamoun, A. W. Rebman, M. E. McClune, E. B. Breitschwerdt, P. J. Reddy, R. Maggi, F. Yang, B. Nemser, A. Ozcan, O. Garner, D. Di Carlo, Z. Ballard, H. A. Joung, A. Garcia-Romeu, R. R. Griffiths, N. Baumgarth, and B. A. Fallon. 2021. Recent Progress in Lyme Disease and Remaining Challenges. Front Med (Lausanne) 8: 666554.

3. Radolf, J. D., K. Strle, J. E. Lemieux, and F. Strle. 2021. Lyme Disease in Humans. Curr Issues Mol Biol 42: 333–384.

4. Steere, A. C., F. Strle, G. P. Wormser, L. T. Hu, J. A. Branda, J. W. Hovius, X. Li, and P. S. Mead. 2016. Lyme borreliosis. Nat Rev Dis Primers 2: 16090.

5. Lochhead, R. B., K. Strle, S. L. Arvikar, J. J. Weis, and A. C. Steere. 2021. Lyme arthritis: linking infection, inflammation and autoimmunity. Nat Rev Rheumatol 17: 449–461.

6. Klempner, M. S., L. T. Hu, J. Evans, C. H. Schmid, G. M. Johnson, R. P. Trevino, D. Norton, L. Levy, D. Wall, J. McCall, M. Kosinski, and A. Weinstein. 2001. Two controlled trials of antibiotic treatment in patients with persistent symptoms and a history of Lyme disease. N Engl J Med 345: 85–92.

7. Marques, A. 2008. Chronic Lyme disease: a review. Infect Dis Clin North Am 22: 341–360, vii-viii.

8. Aucott, J. N., L. A. Crowder, and K. B. Kortte. 2013. Development of a foundation for a case definition of post-treatment Lyme disease syndrome. Int J Infect Dis 17: e443–449.

9. Aucott, J. N., T. Yang, I. Yoon, D. Powell, S. A. Geller, and A. W. Rebman. 2022. Risk of post-treatment Lyme disease in patients with ideally-treated early Lyme disease: A prospective cohort study. Int J Infect Dis 116: 230–237.

10. Chandra, A., G. P. Wormser, A. R. Marques, N. Latov, and A. Alaedini. 2011. Anti-Borrelia burgdorferi antibody profile in post-Lyme disease syndrome. Clin Vaccine Immunol 18: 767–771.

11. Blum, L. K., J. Z. Adamska, D. S. Martin, A. W. Rebman, S. E. Elliott, R. R. L. Cao, M. E. Embers, J. N. Aucott, M. J. Soloski, and W. H. Robinson. 2018. Robust B Cell Responses Predict Rapid Resolution of Lyme Disease. Front Immunol 9: 1634.

12. Bockenstedt, L. K., R. M. Wooten, and N. Baumgarth. 2021. Immune Response to Borrelia: Lessons from Lyme Disease Spirochetes. Curr Issues Mol Biol 42: 145–190.

13. Embers, M. E., N. R. Hasenkampf, M. B. Jacobs, and M. T. Philipp. 2012. Dynamic longitudinal antibody responses during Borrelia burgdorferi infection and antibiotic treatment of rhesus macaques. Clin Vaccine Immunol 19: 1218–1226.

14. Jiang, R., H. Meng, K. Raddassi, I. Fleming, K. B. Hoehn, K. R. Dardick, A. A. Belperron, R. R. Montgomery, A. K. Shalek, D. A. Hafler, S. H. Kleinstein, and L. K. Bockenstedt. 2021. Single-cell immunophenotyping of the skin lesion erythema migrans identifies IgM memory B cells. JCI Insight 6.

15. Barbour, A. G., A. Jasinskas, M. A. Kayala, D. H. Davies, A. C. Steere, P. Baldi, and P. L. Felgner. 2008. A genome-wide proteome array reveals a limited set of immunogens in natural infections of humans and white-footed mice with Borrelia burgdorferi. Infect Immun 76: 3374–3389.

16. Tilly, K., J. G. Krum, A. Bestor, M. W. Jewett, D. Grimm, D. Bueschel, R. Byram, D. Dorward, M. J. Vanraden, P. Stewart, and P. Rosa. 2006. Borrelia burgdorferi OspC protein required exclusively in a crucial early stage of mammalian infection. Infect Immun 74: 3554–3564.

17. Pal, U., X. Yang, M. Chen, L. K. Bockenstedt, J. F. Anderson, R. A. Flavell, M. V. Norgard, and E. Fikrig. 2004. OspC facilitates Borrelia burgdorferi invasion of Ixodes scapularis salivary glands. J Clin Invest 113: 220–230.

18. Tilly, K., A. Bestor, M. W. Jewett, and P. Rosa. 2007. Rapid clearance of Lyme disease spirochetes lacking OspC from skin. Infect Immun 75: 1517–1519.

19. Earnhart, C. G., D. V. Leblanc, K. E. Alix, D. C. Desrosiers, J. D. Radolf, and R. T. Marconi. 2010. Identification of residues within ligand-binding domain 1 (LBD1) of the Borrelia burgdorferi OspC protein required for function in the mammalian environment. Mol Microbiol 76: 393–408.

20. Caine, J. A., and J. Coburn. 2016. Multifunctional and Redundant Roles of Borrelia burgdorferi Outer Surface Proteins in Tissue Adhesion, Colonization, and Complement Evasion. Front Immunol 7: 442.

21. Caine, J. A., Y. P. Lin, J. R. Kessler, H. Sato, J. M. Leong, and J. Coburn. 2017. Borrelia burgdorferi outer surface protein C (OspC) binds complement component C4b and confers bloodstream survival. Cell Microbiol 19.

22. Lin, Y. P., X. Tan, J. A. Caine, M. Castellanos, G. Chaconas, J. Coburn, and J. M. Leong. 2020. Strain-specific joint invasion and colonization by Lyme disease spirochetes is promoted by outer surface protein C. PLoS Pathog 16: e1008516.

23. Tan, X., M. Castellanos, and G. Chaconas. 2023. Choreography of Lyme Disease Spirochete Adhesins To Promote Vascular Escape. Microbiol Spectr 11: e0125423.

24. Branda, J. A., and A. C. Steere. 2021. Laboratory Diagnosis of Lyme Borreliosis. Clin Microbiol Rev 34.

25. Hahm, J. B., J. W. t. Breneman, J. Liu, S. Rabkina, W. Zheng, S. Zhou, R. P. Walker, and R. Kaul. 2020. A Fully Automated Multiplex Assay for Diagnosis of Lyme Disease with High Specificity and Improved Early Sensitivity. J Clin Microbiol 58.

26. Bacon, R. M., B. J. Biggerstaff, M. E. Schriefer, R. D. Gilmore, Jr., M. T. Philipp, A. C. Steere, G. P. Wormser, A. R. Marques, and B. J. Johnson. 2003. Serodiagnosis of Lyme disease by kinetic enzyme-linked immunosorbent assay using recombinant VlsE1 or peptide antigens of Borrelia burgdorferi compared with 2-tiered testing using whole-cell lysates. J Infect Dis 187: 1187–1199.

27. Arnaboldi, P. M., R. Seedarnee, M. Sambir, S. M. Callister, J. A. Imparato, and R. J. Dattwyler. 2013. Outer surface protein C peptide derived from Borrelia burgdorferi sensu stricto as a target for serodiagnosis of early lyme disease. Clin Vaccine Immunol 20: 474–481.

28. Baarsma, M. E., A. Vrijlandt, J. Ursinus, H. L. Zaaijer, S. Jurriaans, A. P. van Dam, and J. W. Hovius. 2022. Diagnostic performance of the ZEUS Borrelia VlsE1/pepC10 assay in European LB patients: a case-control study. Eur J Clin Microbiol Infect Dis 41: 387–393.

29. Coleman, A. S., E. Rossmann, X. Yang, H. Song, C. M. Lamichhane, R. Iyer, I. Schwartz, and U. Pal. 2011. BBK07 immunodominant peptides as serodiagnostic markers of Lyme disease. Clin Vaccine Immunol 18: 406–413.

30. Mathiesen, M. J., M. Christiansen, K. Hansen, A. Holm, E. Asbrink, and M. Theisen. 1998. Peptide-based OspC enzyme-linked immunosorbent assay for serodiagnosis of Lyme borreliosis. J Clin Microbiol 36: 3474–3479.

31. O’Bier, N. S., A. L. Hatke, A. C. Camire, and R. T. Marconi. 2021. Human and Veterinary Vaccines for Lyme Disease. Curr Issues Mol Biol 42: 191–222.

32. Wilske, B., V. Preac-Mursic, S. Jauris, A. Hofmann, I. Pradel, E. Soutschek, E. Schwab, G. Will, and G. Wanner. 1993. Immunological and molecular polymorphisms of OspC, an immunodominant major outer surface protein of Borrelia burgdorferi. Infect Immun 61: 2182–2191.

33. Barbour, A. G., and B. Travinsky. 2010. Evolution and distribution of the ospC Gene, a transferable serotype determinant of Borrelia burgdorferi. MBio 1.

34. Cerar, T., F. Strle, D. Stupica, E. Ruzic-Sabljic, G. McHugh, A. C. Steere, and K. Strle. 2016. Differences in Genotype, Clinical Features, and Inflammatory Potential of Borrelia burgdorferi sensu stricto Strains from Europe and the United States. Emerg Infect Dis 22: 818–827.

35. Seinost, G., D. E. Dykhuizen, R. J. Dattwyler, W. T. Golde, J. J. Dunn, I. N. Wang, G. P. Wormser, M. E. Schriefer, and B. J. Luft. 1999. Four clones of Borrelia burgdorferi sensu stricto cause invasive infection in humans. Infect Immun 67: 3518–3524.

36. Earnhart, C. G., E. L. Buckles, J. S. Dumler, and R. T. Marconi. 2005. Demonstration of OspC type diversity in invasive human lyme disease isolates and identification of previously uncharacterized epitopes that define the specificity of the OspC murine antibody response. Infect Immun 73: 7869–7877.

37. Hanincova, K., P. Mukherjee, N. H. Ogden, G. Margos, G. P. Wormser, K. D. Reed, J. K. Meece, M. F. Vandermause, and I. Schwartz. 2013. Multilocus sequence typing of Borrelia burgdorferi suggests existence of lineages with differential pathogenic properties in humans. PLoS One 8: e73066.

38. Fraser, C. M., S. Casjens, W. M. Huang, G. G. Sutton, R. Clayton, R. Lathigra, O. White, K. A. Ketchum, R. Dodson, E. K. Hickey, M. Gwinn, B. Dougherty, J. F. Tomb, R. D. Fleischmann, D. Richardson, J. Peterson, A. R. Kerlavage, J. Quackenbush, S. Salzberg, M. Hanson, R. van Vugt, N. Palmer, M. D. Adams, J. Gocayne, J. Weidman, T. Utterback, L. Watthey, L. McDonald, P. Artiach, C. Bowman, S. Garland, C. Fuji, M. D. Cotton, K. Horst, K. Roberts, B. Hatch, H. O. Smith, and J. C. Venter. 1997. Genomic sequence of a Lyme disease spirochaete, Borrelia burgdorferi. Nature 390: 580–586.

39. Anderson, J. F., R. A. Flavell, L. A. Magnarelli, S. W. Barthold, F. S. Kantor, R. Wallich, D. H. Persing, D. Mathiesen, and E. Fikrig. 1996. Novel Borrelia burgdorferi isolates from Ixodes scapularis and Ixodes dentatus ticks feeding on humans. J Clin Microbiol 34: 524–529.

40. Steere, A. C., R. L. Grodzicki, A. N. Kornblatt, J. E. Craft, A. G. Barbour, W. Burgdorfer, G. P. Schmid, E. Johnson, and S. E. Malawista. 1983. The spirochetal etiology of Lyme disease. N Engl J Med 308: 733–740.

41. Rudolph, M. J., S. A. Davis, H. M. E. Haque, D. D. Weis, D. J. Vance, C. L. Piazza, M. Ejemel, L. Cavacini, Y. Wang, M. L. Mbow, R. D. Gilmore, and N. J. Mantis. 2023. Structural Elucidation of a Protective B Cell Epitope on Outer Surface Protein C (OspC) of the Lyme Disease Spirochete, Borreliella burgdorferi. mBio 14: e0298122.

42. Kringelum, J. V., C. Lundegaard, O. Lund, and M. Nielsen. 2012. Reliable B cell epitope predictions: impacts of method development and improved benchmarking. PLoS Comput Biol 8: e1002829.

43. Gomes-Solecki, M. J., L. Meirelles, J. Glass, and R. J. Dattwyler. 2007. Epitope length, genospecies dependency, and serum panel effect in the IR6 enzyme-linked immunosorbent assay for detection of antibodies to Borrelia burgdorferi. Clin Vaccine Immunol 14: 875–879.

44. Nemeth, K. L., E. Yauney, J. M. Rock, R. Bievenue, M. M. Parker, and L. M. Styer. 2023. Use of Self-Collected Dried Blood Spots and a Multiplex Microsphere Immunoassay to Measure IgG Antibody Response to COVID-19 Vaccines. Microbiol Spectr 11: e0133622.

45. Yates, J. L., D. J. Ehrbar, D. T. Hunt, R. C. Girardin, A. P. Dupuis, A. F. Payne, M. Sowizral, S. Varney, K. E. Kulas, V. L. Demarest, K. M. Howard, K. Carson, M. Hales, M. Ejemel, Q. Li, Y. Wang, R. Peredo-Wende, A. Ramani, G. Singh, K. Strle, N. J. Mantis, K. A. McDonough, and W. T. Lee. 2021. Serological Analysis Reveals an Imbalanced IgG Subclass Composition Associated with COVID-19 Disease Severity. Cell Reports Medicine: 100329.

46. R core Team. 2023. R: A language and environment for statistical computing. R Foundation for Statistical Computing,.

47. Wickham, H., and J. Bryan. 2023. Read Excel Files. 1.4.3 ed, CRAN. Import excel files into R. Supports ‘.xls’ via the embedded ‘libxls’ C library <https://github.com/libxls/libxls> and ‘.xlsx’ via the embedded ‘RapidXML’ C++ library <https://rapidxml.sourceforge.net/>. Works on Windows, Mac and Linux without external dependencies.

48. Wickham, H. 2019. Welcome to the Tidyverse. Journal of Open Source Software 4: 1686.

49. Wei, T., V. Simko, M. Levy, Y. Xie, Y. Jin, and J. Zemla. 2017. Package ‘corrplot’. Statistician 56: e24.

50. Kolde, R. 2018. pheatmap: Pretty Heatmaps.

51. Lemieux, J. E., W. Huang, N. Hill, T. Cerar, L. Freimark, S. Hernandez, M. Luban, V. Maraspin, P. Bogovic, K. Ogrinc, E. Ruzic-Sabljic, P. Lapierre, E. Lasek-Nesselquist, N. Singh, R. Iyer, D. Liveris, K. D. Reed, J. M. Leong, J. A. Branda, A. C. Steere, G. P. Wormser, F. Strle, P. C. Sabeti, I. Schwartz, and K. Strle. 2023. Whole genome sequencing of human Borrelia burgdorferi isolates reveals linked blocks of accessory genome elements located on plasmids and associated with human dissemination. PLoS Pathog 19: e1011243.

52. Wormser, G. P., D. Brisson, D. Liveris, K. Hanincova, S. Sandigursky, J. Nowakowski, R. B. Nadelman, S. Ludin, and I. Schwartz. 2008. Borrelia burgdorferi genotype predicts the capacity for hematogenous dissemination during early Lyme disease. J Infect Dis 198: 1358–1364.

53. Dimitrov, J. D., C. Planchais, L. T. Roumenina, T. L. Vassilev, S. V. Kaveri, and S. Lacroix-Desmazes. 2013. Antibody polyreactivity in health and disease: statu variabilis. J Immunol 191: 993–999.

54. Mathiesen, M. J., A. Holm, M. Christiansen, J. Blom, K. Hansen, S. Ostergaard, and M. Theisen. 1998. The dominant epitope of Borrelia garinii outer surface protein C recognized by sera from patients with neuroborreliosis has a surface-exposed conserved structural motif. Infect Immun 66: 4073–4079.

55. Earnhart, C. G., D. V. Rhodes, A. A. Smith, X. Yang, B. Tegels, J. A. Carlyon, U. Pal, and R. T. Marconi. 2014. Assessment of the potential contribution of the highly conserved C-terminal motif (C10) of Borrelia burgdorferi outer surface protein C in transmission and infectivity. Pathog Dis 70: 176–184.

56. Dolange, V., S. Simon, and N. Morel. 2021. Detection of Borrelia burgdorferi antigens in tissues and plasma during early infection in a mouse model. Sci Rep 11: 17368.

57. Jobe, D. A., S. D. Lovrich, R. F. Schell, and S. M. Callister. 2003. C-terminal region of outer surface protein C binds borreliacidal antibodies in sera from patients with Lyme disease. Clin Diagn Lab Immunol 10: 573–578.

58. Lovrich, S. D., D. A. Jobe, R. F. Schell, and S. M. Callister. 2005. Borreliacidal OspC antibodies specific for a highly conserved epitope are immunodominant in human lyme disease and do not occur in mice or hamsters. Clin Diagn Lab Immunol 12: 746–751.

59. Izac, J. R., A. C. Camire, C. G. Earnhart, M. E. Embers, R. A. Funk, E. B. Breitschwerdt, and R. T. Marconi. 2019. Analysis of the antigenic determinants of the OspC protein of the Lyme disease spirochetes: Evidence that the C10 motif is not immunodominant or required to elicit bactericidal antibody responses. Vaccine 37: 2401–2407.

60. Jacek, E., K. S. Tang, L. Komorowski, M. Ajamian, C. Probst, B. Stevenson, G. P. Wormser, A. R. Marques, and A. Alaedini. 2016. Epitope-Specific Evolution of Human B Cell Responses to Borrelia burgdorferi VlsE Protein from Early to Late Stages of Lyme Disease. J Immunol 196: 1036–1043.

61. Kirpach, J., A. Colone, J. P. Burckert, W. J. Faison, A. Dubois, R. Sinner, A. L. Reye, and C. P. Muller. 2019. Detection of a Low Level and Heterogeneous B Cell Immune Response in Peripheral Blood of Acute Borreliosis Patients With High Throughput Sequencing. Front Immunol 10: 1105.

62. Hoie, M. H., F. S. Gade, J. M. Johansen, C. Wurtzen, O. Winther, M. Nielsen, and P. Marcatili. 2024. DiscoTope-3.0: improved B-cell epitope prediction using inverse folding latent representations. Front Immunol 15: 1322712.

63. Buckles, E. L., C. G. Earnhart, and R. T. Marconi. 2006. Analysis of antibody response in humans to the type A OspC loop 5 domain and assessment of the potential utility of the loop 5 epitope in Lyme disease vaccine development. Clin Vaccine Immunol 13: 1162–1165.

64. Yang, X., Y. Li, J. J. Dunn, and B. J. Luft. 2006. Characterization of a unique borreliacidal epitope on the outer surface protein C of Borrelia burgdorferi. FEMS Immunol Med Microbiol 48: 64–74.

65. Rebman, A. W., L. A. Crowder, A. Kirkpatrick, and J. N. Aucott. 2015. Characteristics of seroconversion and implications for diagnosis of post-treatment Lyme disease syndrome: acute and convalescent serology among a prospective cohort of early Lyme disease patients. Clin Rheumatol 34: 585–589.

66. Reifert, J., K. Kamath, J. Bozekowski, E. Lis, E. J. Horn, D. Granger, E. S. Theel, J. Shon, J. R. Sawyer, and P. S. Daugherty. 2021. Serum Epitope Repertoire Analysis Enables Early Detection of Lyme Disease with Improved Sensitivity in an Expandable Multiplex Format. J Clin Microbiol 59.

67. Cobaugh, C. W., J. C. Almagro, M. Pogson, B. Iverson, and G. Georgiou. 2008. Synthetic antibody libraries focused towards peptide ligands. J Mol Biol 378: 622–633.

68. Blum, J. S., M. L. Fiani, and P. D. Stahl. 1991. Proteolytic cleavage of ricin A chain in endosomal vesicles. Evidence for the action of endosomal proteases at both neutral and acidic pH. J Biol Chem 266: 22091–22095.

69. Trier, N. H., and G. Houen. 2023. Antibody Cross-Reactivity in Auto-Immune Diseases. Int J Mol Sci 24.

70. Boughter, C. T., M. T. Borowska, J. J. Guthmiller, A. Bendelac, P. C. Wilson, B. Roux, and E. J. Adams. 2020. Biochemical patterns of antibody polyreactivity revealed through a bioinformatics-based analysis of CDR loops. Elife 9.

71. Mouquet, H., J. F. Scheid, M. J. Zoller, M. Krogsgaard, R. G. Ott, S. Shukair, M. N. Artyomov, J. Pietzsch, M. Connors, F. Pereyra, B. D. Walker, D. D. Ho, P. C. Wilson, M. S. Seaman, H. N. Eisen, A. K. Chakraborty, T. J. Hope, J. V. Ravetch, H. Wardemann, and M. C. Nussenzweig. 2010. Polyreactivity increases the apparent affinity of anti-HIV antibodies by heteroligation. Nature 467: 591–595.

72. Ikushima, M., F. Yamada, S. Kawahashi, Y. Okuyama, and K. Matsui. 1999. Antibody response to OspC-I synthetic peptide derived from outer surface protein C of Borrelia burgdorferi in sera from Japanese forestry workers. Epidemiol Infect 122: 429–433.

73. Pulzova, L., Z. Flachbartova, E. Bencurova, L. Potocnakova, L. Comor, E. Schreterova, and M. Bhide. 2016. Identification of B-cell epitopes of Borrelia burgdorferi outer surface protein C by screening a phage-displayed gene fragment library. Microbiol Immunol 60: 669–677.

74. Tokarz, R., N. Mishra, T. Tagliafierro, S. Sameroff, A. Caciula, L. Chauhan, J. Patel, E. Sullivan, A. Gucwa, B. Fallon, M. Golightly, C. Molins, M. Schriefer, A. Marques, T. Briese, and W. I. Lipkin. 2018. A multiplex serologic platform for diagnosis of tick-borne diseases. Sci Rep 8: 3158.

75. Yu, Z., J. M. Carter, L. H. Sigal, and S. Stein. 1996. Multi-well ELISA based on independent peptide antigens for antibody capture. Application to Lyme disease serodiagnosis. J Immunol Methods 198: 25–33.

76. Baum, E., A. Z. Randall, M. Zeller, and A. G. Barbour. 2013. Inferring epitopes of a polymorphic antigen amidst broadly cross-reactive antibodies using protein microarrays: a study of OspC proteins of Borrelia burgdorferi. PLoS One 8: e67445.

