## Supplemental Tables 1-2 and Figure for "A refined human linear B cell epitope map of Outer surface protein C (OspC) from the Lyme disease spirochete, *Borreliella burgdorferi*"

**Table S1: IgG reactivity (>3SD) to $OspC_{A/B/K}$ and OspC peptides in diagnostic and PTLD serum samples**

| | Diagnostic | | PTLD | |
|---|---|---|---|---|
| **OspC / peptide** | **n (%)[a]** | **Fold increase (SD)[b]** | **n (%)[a]** | **Fold increase (SD)[b]** |
| $OspC_A$ | 240 (34.5) | 3.7 (3.2) | 43 (27.2) | 7.0 (9.0) |
| $OspC_B$ | 388 (55.7) | 5.0 (5.4) | 6 (3.8) | 3.6 (1.3) |
| $OspC_K$ | 243 (34.9) | 6.1 (6.4) | 38 (24.1) | 3.9 (3.7) |
| $9\text{-}33_{ABK}$ | 7 (1.0) | 1.8 (1.2) | 1 (0.6) | 1.2 (N/A) |
| $40\text{-}55_{AB}$ | 45 (6.5) | 2.3 (2.0) | 7 (4.4) | 2.3 (1.1) |
| $40\text{-}55_{K}$ | 16 (2.3) | 1.9 (1.0) | 5 (3.2) | 1.8 (0.6) |
| $71\text{-}86_{A}$ | 92 (13.2) | 1.7 (1.1) | 5 (3.2) | 1.5 (0.5) |
| $132\text{-}146_{A}$ | 46 (6.6) | 2.5 (2.2) | 9 (5.7) | 2.4 (1.3) |
| $132\text{-}146_{B}$ | 118 (17.0) | 2.1 (1.8) | 21 (13.3) | 3.2 (2.9) |
| $133\text{-}147_{K}$ | 75 (10.8) | 2.6 (3.3) | 15 (9.5) | 2.9 (2.1) |
| $155\text{-}169_{A}$ | 75 (10.8) | 2.6 (3.3) | 10 (6.3) | 2.8 (2.4) |
| $177\text{-}186_{A}$ | 82 (11.8) | 1.9 (1.0) | 28 (17.7) | 2.7 (2.7) |
| $178\text{-}187_{K}$ | 164 (23.6) | 1.8 (1.0) | 30 (19.0) | 3.1 (3.7) |
| $178\text{-}187_{B}$ | 24 (3.4) | 1.5 (0.4) | 25 (15.8) | 2.7 (2.7) |
| $183\text{-}190_{AB}$[c] | 110 (15.8) | 2.1 (1.4) | 22 (13.9) | 3.7 (4.5) |
| $193\text{-}210_{A}$ (C10) | 272 (39.1) | 5.4 (6.8) | 36 (22.8) | 3.8 (4.3) |
| VlsE C6-17 | 537 (77.2) | 9.2 (5.3) | 78 (49.4) | 7.8 (8.1) |

[a], The number of samples (n =) and percent reactivity ("%") to each antigen in the diagnostic (n=696) and PTLD (n=158) sample sets were calculated using a cutoff of >3SD above the mean of controls samples; [b], The fold increase of positive sample antibody reactivity over controls such than an index value of 1 is 3SD above the control mean, while 3.7 corresponds to ~11SD greater than the mean control. [c], $OspC_B$ residues 184-191

**Table S2: Serum IgM reactivity greater than 3SD and 6SD to $OspC_{A/B/K}$ and OspC peptides in diagnostic serum samples**

| OspC / peptide | 3SD n = (%)[a] | 6SD n = (%)[a] |
|---|---|---|
| $OspC_A$ | 398 (57.2) | 306 (44.0) |
| $OspC_B$ | 309 (44.4) | 234 (33.6) |
| $OspC_K$ | 219 (31.5) | 172 (24.7) |
| $9\text{-}33_{ABK}$ | 22 (3.2) | 5 (0.7) |
| $40\text{-}55_{AB}$ | 25 (3.6) | 1 (0.1) |
| $40\text{-}55_{K}$ | 18 (2.6) | 5 (0.7) |
| $71\text{-}86_{A}$ | 24 (3.4) | 4 (0.6) |
| $132\text{-}146_{A}$ | 32 (4.6) | 9 (1.3) |
| $132\text{-}146_{B}$ | 32 (4.6) | 4 (0.6) |
| $133\text{-}147_{K}$ | 38 (5.5) | 6 (0.9) |
| $155\text{-}169_{A}$ | 57 (8.2) | 19 (2.7) |
| $177\text{-}186_{A}$ | 27 (3.9) | 1 (0.1) |
| $178\text{-}187_{K}$ | 170 (24.4) | 61 (8.8) |
| $178\text{-}187_{B}$ | 100 (14.4) | 40 (5.7) |
| $183\text{-}190_{AB}$[b] | 32 (4.6) | 5 (0.7) |
| $193\text{-}210_{A}$ (C10) | 57 (8.2) | 14 (2.0) |

[a], The number of samples (n =) and percent reactivity ("%") to each antigen in the diagnostic (n=696) sample set were calculated using cutoffs of >3SD and >6SD above the mean of controls samples; [b], Residues 184-191 in Type B.

**Figure S1**

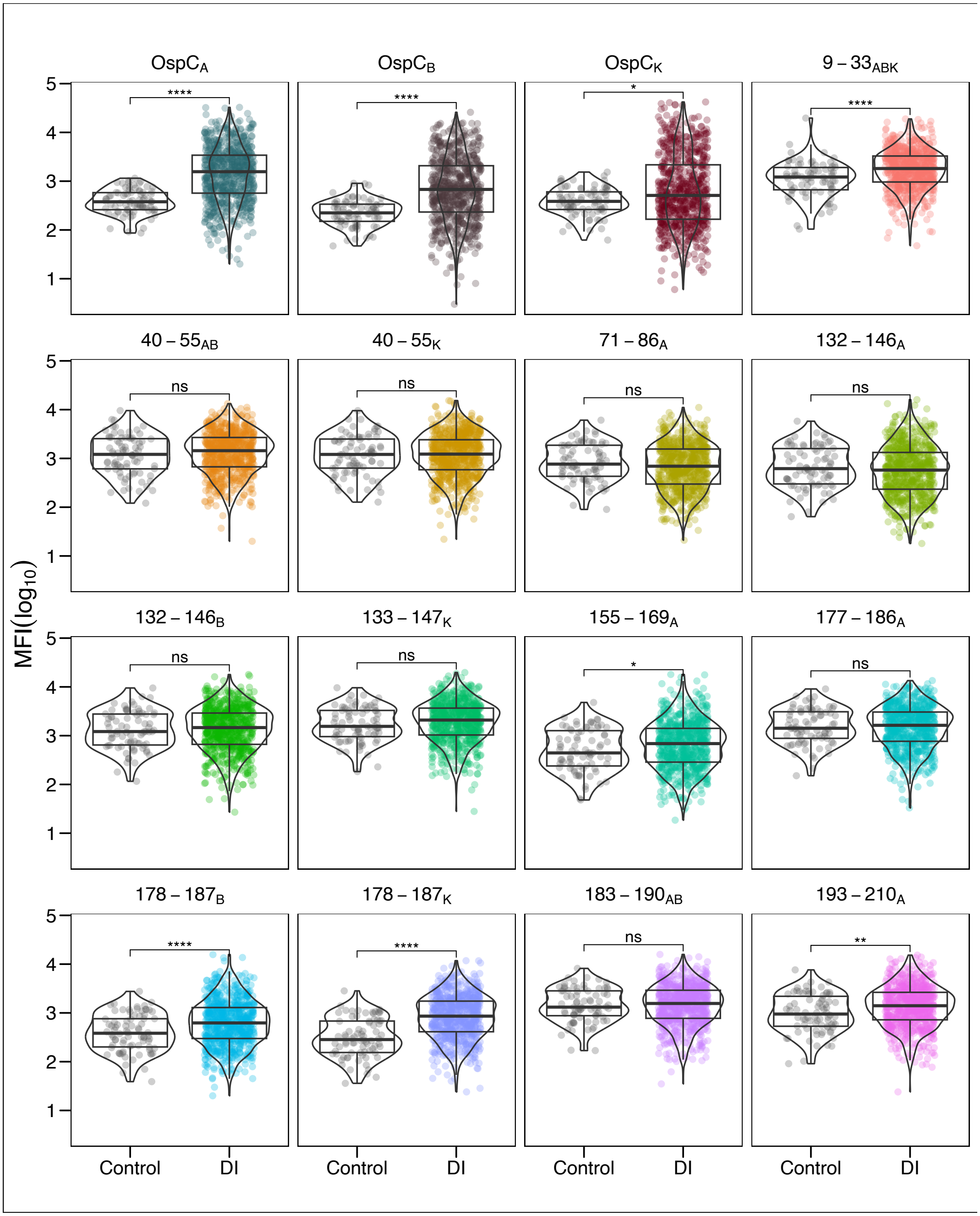


**Figure S1. Serum IgM reactivity of diagnostic samples with OspC and OspC-derived peptides.** DI serum samples (diluted 1:100) were subject to MIA using microspheres coated with recombinant dimeric OspC types A, B and K and OspC-peptides as described in Table 1. Panels are labeled by corresponding residue numbers for each OspC type. MFI values were $\log_{10}$ transformed and compared to the control sample set. Significance was determined by the Mann-Whitney U-test and Student t-test depending on data distribution (*, $p < 0.05$).
